# Resting-state connectivity predicts patient-specific effects of deep brain stimulation for Parkinson’s disease

**DOI:** 10.1101/203406

**Authors:** Xiaoyu Chen, Chencheng Zhang, Yuxin Li, Pei Huang, Qian Lv, Wenwen Yu, Shengdi Chen, Bomin Sun, Zheng Wang

## Abstract

Neural circuit-based guidance for optimizing patient screening, target selection and parameter tuning for deep brain stimulation (DBS) remains limited. To this end, we propose a functional brain connectome-based modeling approach that simulates network-spreading effects of stimulating different brain regions and quantifies rectification of abnormal network topology *in silico*. We validate these analyses by predicting nuclei in basal-ganglia circuits as top-ranked targets for 43 local patients with Parkinson’s disease and 90 patients from public database. However, individual connectome-based predictions demonstrate that globus pallidus and subthalamic nucleus (STN) constituted as the best choice for 21.1% and 19.5% of patients, respectively. Notably, the priority rank of STN significantly correlated with motor symptom severity in the local cohort. By introducing whole-brain network diffusion dynamics, these findings unfold a new dimension of brain connectomics and underscore the importance of neural network modeling for personalized DBS therapy, which warrants experimental investigation to validate its clinical utility.

---

Brain stimulation via optic, sonic, electrical and magnetic means enables the adjustable and selectable modulation of network-level activity, thereby producing varying degrees of therapeutic effects. As one of the most successful neuromodulation-based clinical interventions, deep brain stimulation (DBS) has demonstrated remarkable symptomatic amelioration in a wide range of neurological ^1, 2, 3, 4, 5^, and psychiatric conditions ^6, 7^. However, the clinical efficacy of brain stimulation is unpredictable on a case-by-case basis, as it is inevitably susceptible to unexpected postoperative side effects on cognitive function, mood and behavior ^8, 9, 10^. This may be partially caused by our incomplete understanding of its neural circuit-level mechanisms ^2, 4, 5, 11, 12^. Meanwhile, considerable variability in terms of heterogeneous neuropathology, clinical trajectory and treatment protocols exists between individual patients. To date, there are no theoretical principles or pre-surgical consensuses for the determination of desirable stimulation targets (e.g. pallidal versus subthalamic DBS)^13, 14, 15, 16, 17^ and the fine-tuning of stimulation parameters (e.g., current amplitude, frequency, and pulse-width etc.)^4, 9, 10^ to optimize a single patient’s outcome. Inter-individual differences in stimulation-induced effects therefore pose a tremendous challenge for pertinent empirical studies and therapeutic interventions alike, heightening the urgent demand for optimizing patient-specific stimulation protocols ^15, 18^.

The functional brain connectome constructed from resting-state functional connectivity is powerful in characterizing the nature of topological organization ^19, 20, 21, 22, 23^ and the neuropathology of the diseased brain ^24, 25, 26, 27^. Grounded in network science and graph theory, the human connectome can be readily formulated as a brain graph or matrix consisting of nodes (parcellated brain regions) and edges (connections between nodes)^20^. Emerging evidence suggests that both invasive and noninvasive therapeutics can reconfigure brain networks and normalize maladaptive functional connectivity of brain circuitry concomitantly with clinical symptomatic improvement ^28, 29, 30^. In other words, the therapeutic effects of neurostimulation are attributable to the rectification or rebalance of abnormal network topology associated with pathological behaviors ^11, 31, 32, 33^. This has enabled the development of a variety of whole-brain computational models with an emphasis on clinical applications ^18, 19, 25, 26, 34^. Nevertheless, a majority of current fMRI studies employ the case-control approach to detect group effects by leveraging the statistical benefits of averaging across subjects. They typically ignore the considerable heterogeneity of clinical cohorts and draw inferences about general patterns of brain pathology commonly shared across a cluster of patients, which is not directly applicable to clinical situations. A growing number of studies have demonstrated that an individual brain can be differentially characterized by its connectome in both healthy ^35, 36, 37^ and diseased conditions ^38, 39^. However, no existing neural circuit-based methods can be harnessed to guide patient screening or to predict treatment outcome before surgical intervention ^2, 4, 31^.

To address these bedrock issues, we propose a functional connectome-based neuromodeling approach by which we are able to predict neurostimulation targets and strengths at both population and individual levels (as illustrated in Figure 1). The basic intuition is that quantitative analysis of the rectification of dysfunctional connectivity towards a healthy regime in an individual patient at the brain-wide scale gives rise to the optimal determination of stimulation conditions. In comparison to a group of 46 healthy subjects, we launched a proof-of-principle test in two patient groups: 43 PD patients recruited from the local community and 90 PD patients from a public dataset (as an independent validation). Using fully cross-validated analysis, we identified brain areas mainly located in basal-ganglia circuits including the globus pallidus and subthalamic nucleus, as prime targets in 86.1% of the local and 87.8% of the public cohort. We further determined the optimal range of neurostimulation strength for each targeted area. This computational study allows the exploration of a wide array of potential therapeutic brain regions beyond basal-ganglia circuits, and guidance of fine-tuning stimulation parameters in individual patients with Parkinson’s disease, which can be integrated as part of a pre-surgical treatment plan in future trials. Although thus far only substantiated using PD data, it can be readily generalized to other neuropsychiatric disorders and therapeutic stimulation modalities (e.g., transcranial magnetic stimulation), providing a general analytics whole-brain modeling framework for brain stimulation at the macroscale.

**Figure 1.**
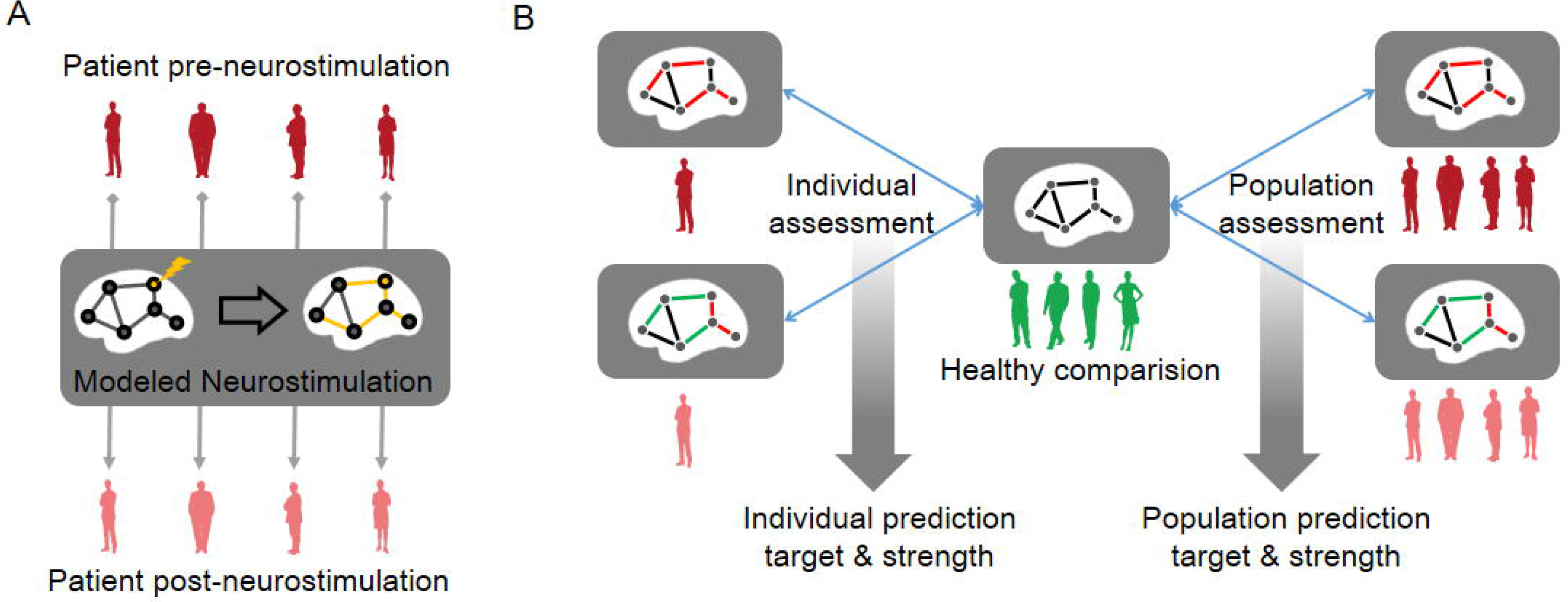
A schematic diagram of the whole-brain neurostimulation modeling in both population and individual levels. **(A)** The pre-neurostimulation functional brain connectome of an individual PD patient (red) is subjected to focal stimulation of a selected target site (yellow node), where neurostimulation generates globally distributed network effects to the whole brain (yellow edges), and then results in a post-neurostimulation connectome (pink). **(B)** Compared to the healthy connectome (middle, green), abnormal network topology in a single patient (left) or patient population (right) may be partially normalized by neurostimulation. The neurostimulation-induced topological rebalance effects can be assessed on both population-averaged and individual connectomes.

## Results

### Internal validation: target and strength prediction

We began by screening all 46 bilateral brain regions as potential targets in the local PD group. The most effective neurostimulation strength for each region (resulting in the largest percentage of improvement) was used to compare outcomes among all regions. At group level, the globus pallidus (GP) was clearly identified as the best choice for neurostimulation, exhibiting the largest relative change of 2.22% (Figure 2A). This indicates that the greatest improvement in the local PD group may be achieved by focal modulation of the GP. The STN, thalamus and putamen evidently emerged as top-ranked targets with relative changes of 0.97%, 1.48% and 1.17%, respectively. The hippocampus was found to be a desirable target for which neurostimulation could attain a 1.00% improvement. Importantly, these results withstood a 1000-round sub-sampling cross-validation process (Figure S1A and S1B). Using a delicate 1024-region parcellation template to repeat the analysis, we confirmed that the GP, STN, putamen, thalamus, and hippocampus remained top-ranked candidates for neurostimulation (Figure S2). Note that the performance of different subregions within the thalamus varies widely, suggesting that some particular nuclei of the thalamus could be extremely sensitive to stimulation. Furthermore, we randomly selected half of the healthy control sample as “patients” for target prediction to demonstrate that this model would not biasedly predict any specific brain region as a potential target (Figure S3A and S3B). Meanwhile, neurostimulation strength, another key element of a typical stimulation protocol, was estimated for each candidate target. In the local PD group, optimal neurostimulation strengths for top-five targets including the GP, STN, putamen, thalamus and hippocampus were determined as 50%, 56%, 36%, 32% and 32% down-regulation of their original connectivity strengths, respectively (Figure 2B). Predicted strengths for the remaining regions are displayed in Figure S4.

**Figure 2.**
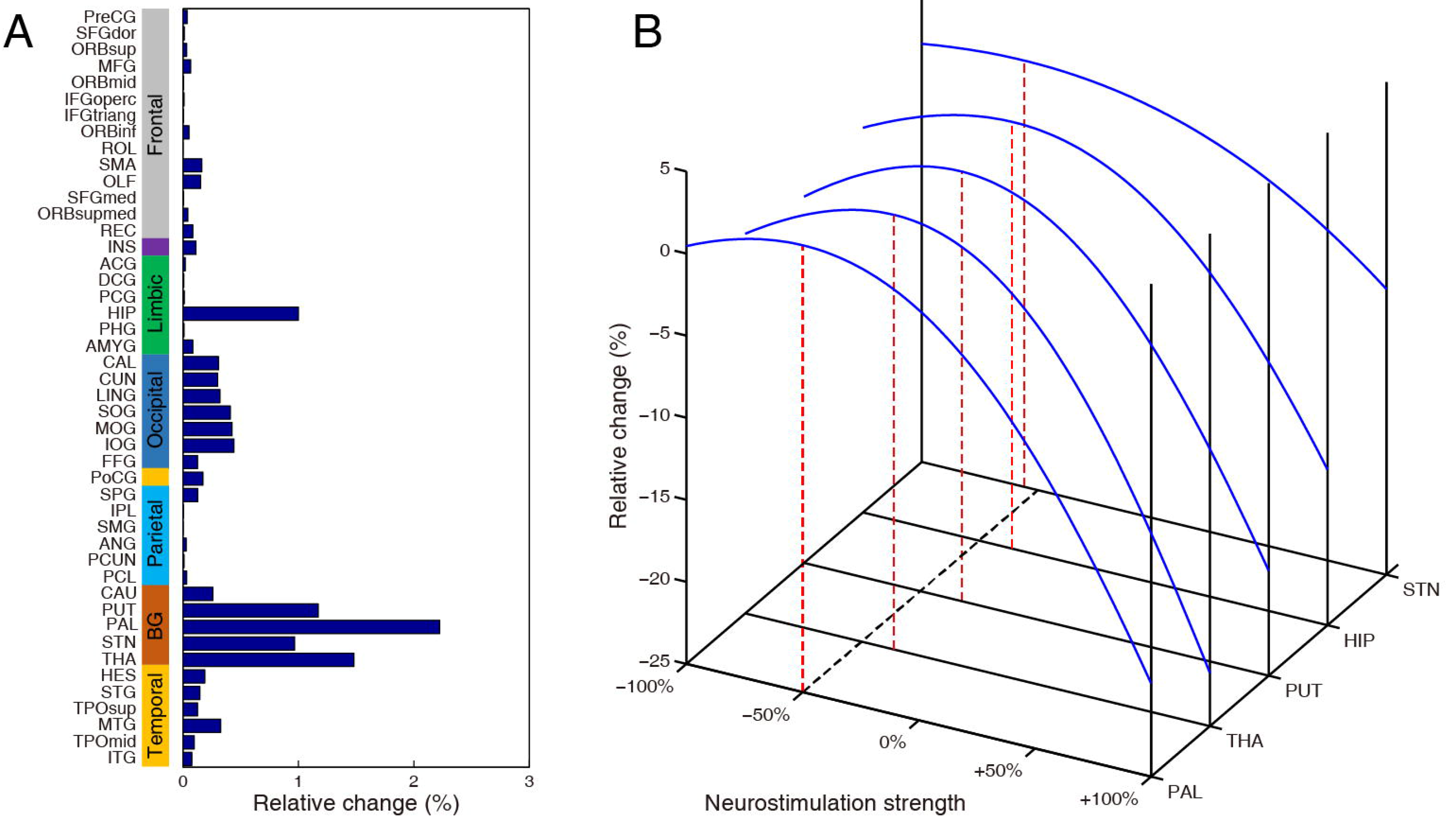
Neurostimulation target prediction for the local PD group. **(A)** The highest relative changes of all brain regions after neurostimulation at the population level are plotted for the local cohort. The parcellated regions are shown on the Y-axis and categorized into frontal, insula, limbic, occipital, parietal, and temporal lobes, and the basal ganglia. **(B)** Relative change is plotted against varying neurostimulation strengths for each top-five target. Red bars mark the optimal neurostimulation strength with the largest relative change at each region.

We then moved on to target and strength prediction for individual patients. For each brain region, we plotted the relative change in individual patients (mean ± SEM, left panel of Figure 3A). In the local cohort, the top-five brain regions, including the GP, STN, thalamus, putamen and hippocampus, consistently demonstrate significant improvement (p < 10^−11^, one sample t-test, indicated by stars in left panel of Figure 3A), with rather large effect sizes (Cohen’s *d* > 1.5, right panel of Figure 3A). We subsequently plotted the complete rank of all regions as potential candidates for each patient in the local cohort (left panel of Figure 3B) and counted their occurring frequencies as the best or top five choice (right panel of Figure 3B). Evidently, the GP and STN are the best choices for 11 (25.6%) and 11 (25.6%) patients (pie plot in the right panel of Figure 3B) and top-five choices for 33 (76.7%) and 36 (83.7%) patients, respectively. Meanwhile, the caudate, thalamus and hippocampus are the best targets for 9 (20.9%), 6 (14.0%), and 4 (9.3%) patients, and top-five choices for 22 (51.2%), 28 (65.1%), and 25 (58.1%) patients, respectively. In sum, the nuclei of the basal ganglia circuit are predicted as the best choices for 86.1% and as top-five choices for 97.7% of the local cohort, upholding striking agreement with prior clinical results ^40^. Intriguingly, we found a significant relationship between the rank of the STN in this cohort and the severity of symptoms as indexed by UPDRS-III (p < 0.05, Kendall rank correlation), but not for any other top sites. Moreover, we made personalized predictions of optimal strengths for each region in individual patients of the local cohort (left panel of Figure 4), where hot colors represent up-regulation and cold colors represent down-regulation of the original connectivity. Regardless of marked variability between individuals, consistent down-regulation (p < 10^−11^, right panel of Figure 4) of basal ganglia- and hippocampal- related functional connectivity substantially contributes to the rectification of dysfunctional networks.

**Figure 3.**
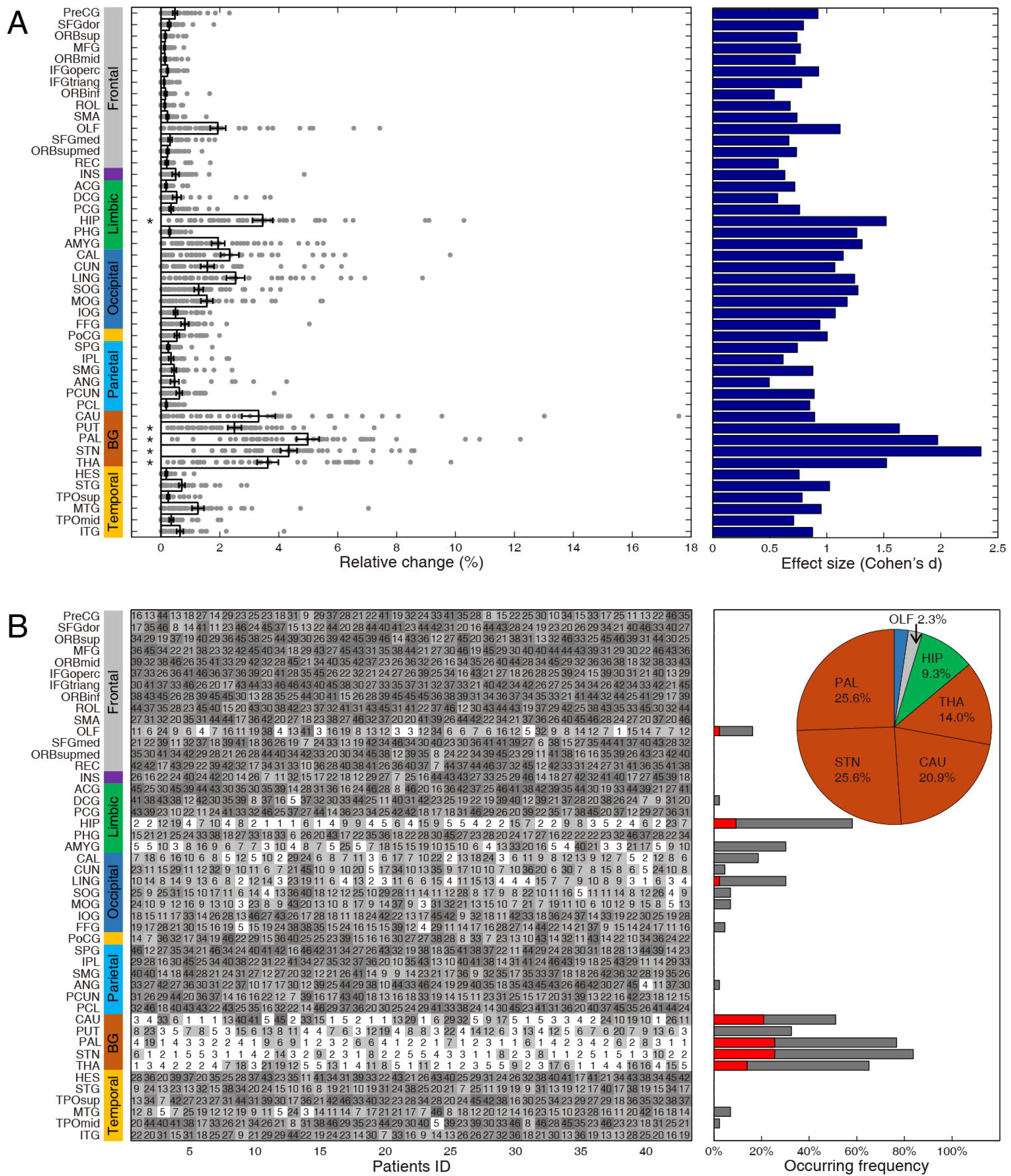
Neurostimulation target prediction for individual local PD patients. **(A)** In the left panel, the relative changes (x-axis) of all brain regions for each patient (denoted as dots) are plotted, in which the bar plot with error bars represents mean ± SEM (* indicates top-five targets with statistical significance p<10^−11^, one sample t-test). The corresponding effect sizes (Cohen’s *d* value) for each brain region are illustrated in the right panel. **(B)** In the left panel, a complete rank of all brain regions is plotted for each patient; the x-axis represents the patient ID in the local PD cohort. Top-five brain regions of each patient are highlighted with white color. In the right panel, the occurring frequencies of the best and top-five sites are counted and displayed as red and grey bars, respectively. The pie plot summarizes the occurring frequency of the best target for each patient of this local group.

**Figure 4.**
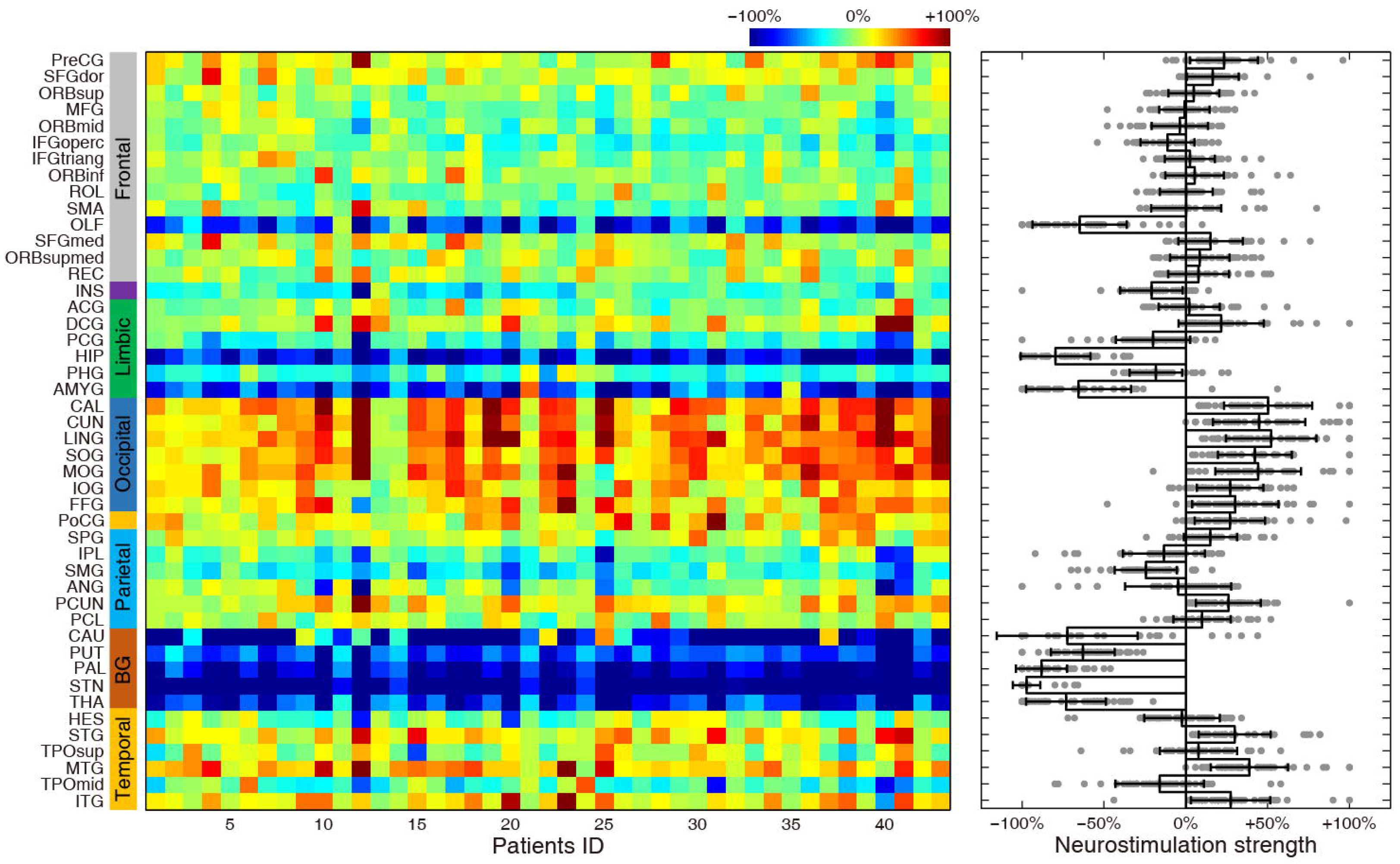
Neurostimulation strength prediction for individual local PD patients. In the left panel, the optimal neurostimulation strengths for all regions for each patient of the local group are illustrated. The color bar represents the range of simulated neurostimulation strengths. In the right panel, the neurostimulation strengths (x-axis) of all regions are plotted for all patients (denoted as dots); the bar plot with error bars represents mean ± SEM.

### External validation: target and strength predictions

As a robust test of generalizability, we applied the present modeling analysis to a completely independent validation dataset from the public PPMI database, which provides resting-state fMRI data for 90 patients with PD (Figure 5). We observed that nuclei of the basal ganglia circuit and hippocampus stand out as the prioritized selections, in agreement with results from the local PD cohort (Figure 5A). Correspondingly, we found that optimal neurostimulation strengths for the GP, STN, caudate, putamen and hippocampus are almost identical to those in the local group (58%, 82%, 84%, 40% and 52% down-regulation shown in Figure 3B, respectively). At the single subject level, the predicted top-five targets for each patient in the public cohort are also consistent with those of the local group. Stimulation of regions including the GP, STN, thalamus, putamen, hippocampus and amygdala demonstrates significant percentage improvement (p<10^−20^, one sample t-test, left panel of Figure 5C) in this group with large effect sizes (Cohen’s *d* > 1.3, right panel of Figure 5C). The GP and STN are the best targets for 17 (18.9%) and 15 (16.7%) patients, and top-five choices for 77 (85.6%) and 67 (74.4%) patients, respectively (Figure 5D). The caudate, thalamus, hippocampus and amygdala are the best choices for 45 (50.0%), 2 (2.2%), 8 (8.9%) and 1 (1.1%) individual patients, respectively, and top-five choices for 66 (73.3%), 50 (55.6%), 75 (83.3%) and 24 (26.7) patients (Figure5D). We can therefore summarize the nuclei of the basal ganglia circuits as prioritized targets for 87.8% and as top-five choices for 100% of this public group. Note that we did not observe a statistically significant relationship between the ranks of any top-five sites and the severity of motor symptoms in the public cohort (p > 0.05, Kendall rank correlation). Meanwhile, personalized predictions of optimal strength for each region are shown for the public cohort in Figure 5E, as manifested consistent down-regulation.

**Figure 5.**
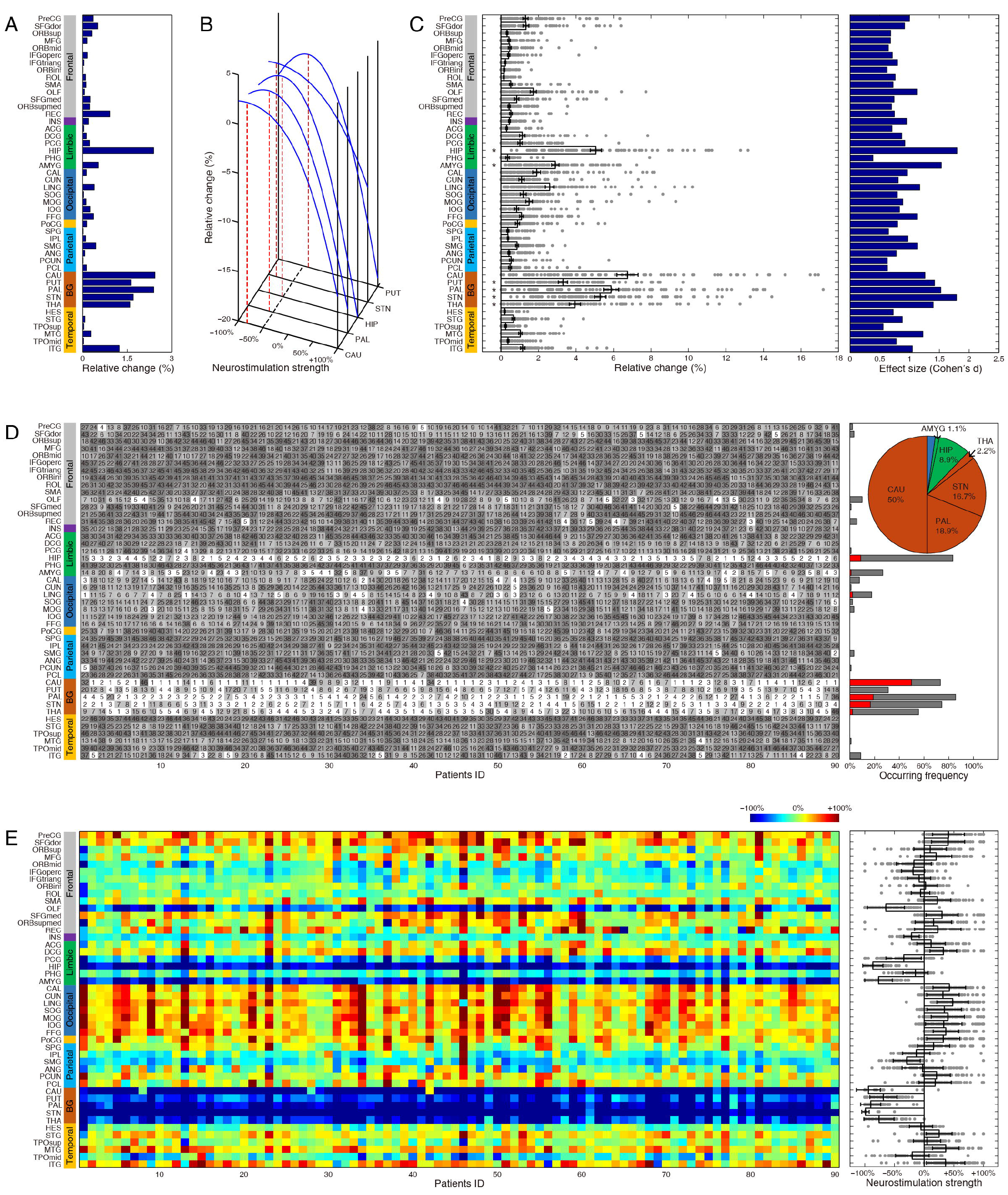
Prediction of neurostimulation target and strength for the public PD cohort. Predictive results of the public group including targets **(A)** and strengths **(B)** are shown in the same manner as Figure 2. Predictive results of individual patients of this public group are shown in (**C-E**). In the left panel of (**C**), each dot stands for one patient, and the bar plot with error bars represents mean ± SEM of the relative change (*indicates top-five targets with statistical significance p<10^−20^, one sample t-test).

### Reconfiguration of network topology by neurostimulation

We investigated whether and how focal stimulation of the GP or STN resulted in topological reconfiguration of the functional brain connectome in local (Figure 6) and public cohorts (Figure S5). The removed (green) and remaining (grey) abnormal connections after neurostimulation are illustrated as compared to the pre-stimulation connectomic matrix (p < 0.05, Bonferroni corrected for multiple comparisons). Neurostimulation remarkably resulted in the removal of a large number of abnormal connections between the target regions and widespread areas in temporal, parietal, frontal and limbic brain regions. Although not found in the local cohort, considerable abnormal connections in the basal ganglia circuits that bypass the GP (Figure S5A and B) or STN (Figure S5E and F) were eliminated in the public cohort. Findings for the public PD cohort were largely comparable, though a substantial number of connections emerged from the intervention process. Taken together, these results demonstrate that focal stimulation leads to remarkable topological reconfiguration toward healthy bifurcation of the functional connectomes as measured in controls, suggesting potential therapeutic mechanisms for the alleviation of clinical symptoms.

**Figure 6.**
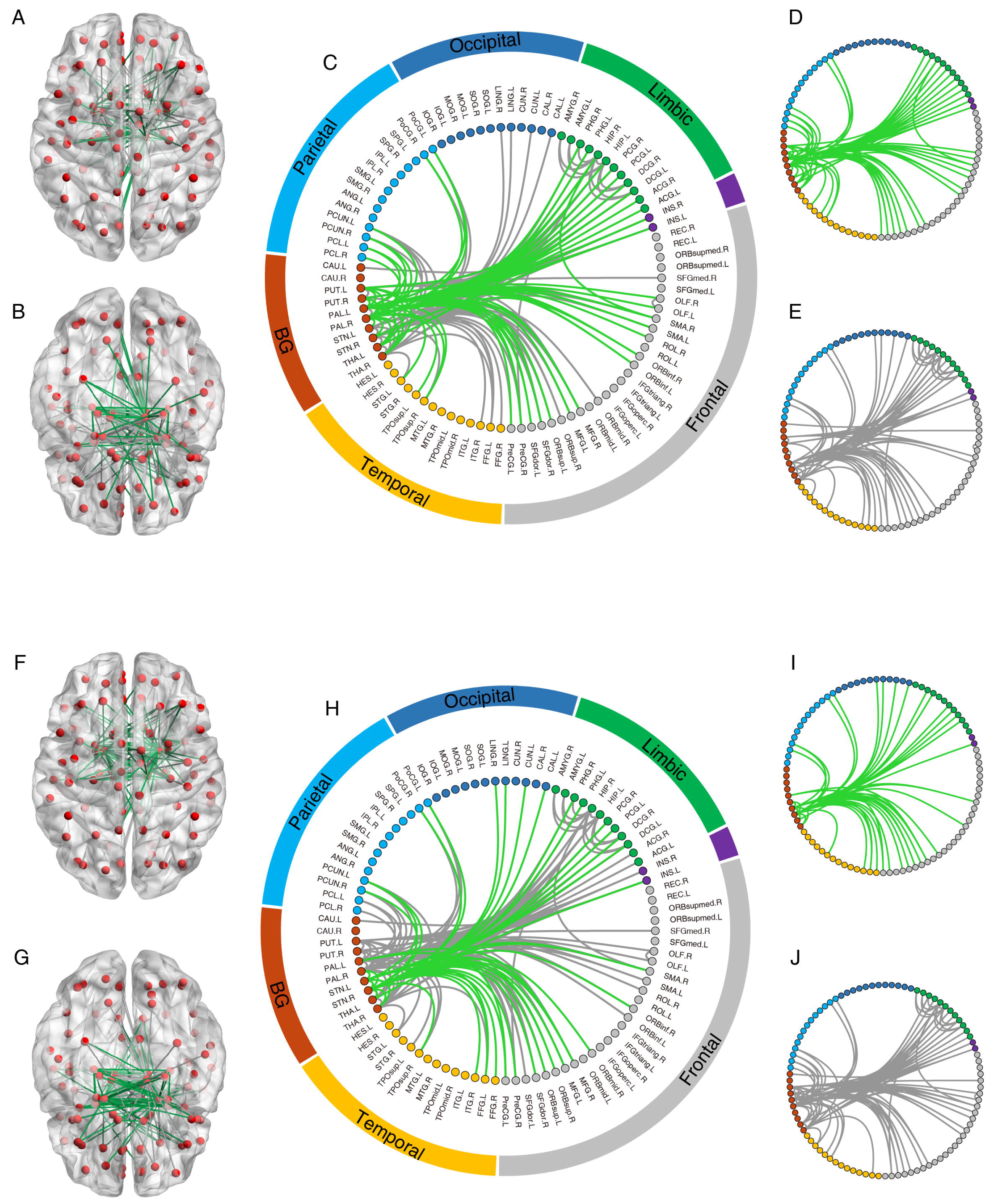
Representative topological changes of the functional brain connectome induced by stimulating GP (**A-E**, top panel) and STN (**F-J**, bottom panel) for the local PD cohort. All involved brain regions (nodes, shown as red spheres) and connections (edges, shown as lines between nodes) were rendered on a transparent brain from superior (**A**) and inferior (**B**) axial view angles. As illustrated in **C**, all connections were categorized into different brain lobes. Green lines represent the removed abnormal connections that were significantly different between the local PD cohort and healthy comparison subject group (p < 0.05 corrected) before, but not after, neurostimulation. Grey lines represent the remaining abnormal connections after neurostimulation (p < 0.05 corrected). (**D-E**) Illustration of altered connections indicated by green and grey lines in (**C**). (**F-J**) Illustration of topological changes of functional connections if stimulating STN for the local PD cohort.

## Discussion

The present whole-brain modeling approach indeed describes a fundamental biological process with an analytic solution, and then translates the process of simulating the global effects of focal stimulation for clinically relevant purposes. It also suggests that spontaneous functional connectivity encoded in brain networks is sufficiently sensitive to capture the predisposition of individual’s susceptibility in Parkinson’s disease. Pioneering work based on the structural brain connectome or in combination with functional brain connectome has shown promising results that model the dynamic network effects of a neural insult (lesion or stimulation), and predict network spreading and functional consequences ^32, 41, 42, 43, 44, 45^. The present approach requires only resting-state functional MRI data of individuals to derive an empirical subject-specific transformation rule from the local to the global scale. This type of network diffusion process may stand for a fundamental process widely applicable to various biological systems, in that aberrant activity easily spreads through connected nodes and induces extensive pathological effusions in the system. Moreover, we posit that reversal of abnormal network topology can be indicative of a positive outcome for brain stimulation therapy *in silico*, which formulates the fundamental basis for the predictions of pre-stimulation protocols in individuals, including candidate screening, target selection and parameter tuning.

Target selection remains an unresolved challenge in the field of deep-brain stimulation in both neurological and psychiatric conditions ^1, 6, 46, 47^. Insights gained from small animal experiments typically involve tedious procedures in which multiple brain regions are sequentially stimulated ^48^, and are not directly applicable as guidance for surgical procedure in human subjects ^49^. Our resting-state connectivity-based approach was able to predict nuclei of the basal ganglia circuits as optimal targets in two independent cohorts, in striking accordance with previous reports and clinical treatment routines for various movement disorders ^3, 13, 14, 15, 40, 50, 51^. Nevertheless, we observed a rather large proportion of patients (51.2% in local cohort, 35.6% in public cohort) for whom the best target was neither the GP nor the STN. In a very few patients (as shown in Figure 3B), all the nuclei of the basal ganglia circuits ranked extremely low, suggesting that focal stimulation therapeutics may not be appropriate for them at all. Hence, our computational approach provides a more tailored strategy to guide individualized selection between these candidate targets, thereby improving the overall outcome of neurostimulation ^15^. It has been shown that the thalamus was the best choice for stimulation in patients with severe tremor-related symptoms ^52^. The putamen, another key component of the basal ganglia circuits, was identified among top-ranking candidates, which is consistent with prior experience in patients ^53^. Interestingly, the caudate emerged as a good candidate for some patients, especially in the public cohort (Figure 5), despite its occurrence as a common target for stimulation in patients with epilepsy ^47^ or psychiatric conditions ^6^. Treatment outcomes for the caudate had high variation as may lead to relatively small effect size. Surprisingly, the hippocampus was identified here as a potential target, most likely because it emerged as one of key abnormal nodes comparing the brain matrices of patients to controls. And it is known to be pathologically involved in the non-motor clinical symptoms of Parkinsonism ^54^. However, it has not been reported hitherto in the clinical treatment of PD patients and future investigation in animal and human subjects is required to test experimental outcomes of stimulation in this area.

Neurostimulation strength is another critical parameter affecting therapeutic outcomes in individuals. Determination of optimal strength in our modeling is analogous to the programming of a DBS device in individual patients, which often demands labor-intensive adjustment based on frequent symptom assessment by clinicians. In the present study, the priority of all brain regions as potential neurostimulation targets is ranked in each single patient when the optimal stimulation strength is applied to each area. Our predictions clearly show that excessively strong or weak neurostimulation may not result in desirable modulation even at an appropriately selected target, and that optimal strengths (including up- or down-regulation) vary substantially among different targets across patients. These findings may serve as theoretical guidance for the tuning of stimulation protocols and possibly obviate the need for testing by trial and error on human subjects. This is a rather attractive premise, as brain stimulation with inappropriate parameters usually induces certain adverse effects and shortens battery life ^2, 15^. More importantly, the present integrated strategy of predicting stimulation targets and strengths can be implemented on the basis of single patients, thereby opening a new avenue for the derivation of personalized treatments in phenomenologically inhomogeneous populations.

At the current stage, this connectome-based computational method has several practical limitations. First, the precise correspondence between metrics of the computational model (e.g. neurostimulation strength) and the parameters of a stimulation device (such as current amplitude and frequency of stimulus pulse) merits future *in vivo* experimental validation. The device must be calibrated so that the actual dose of stimulation is known. Future retrospective or prospective clinical investigation in patients with DBS implantation would invaluably facilitate this translational application. Second, the prediction accuracy of the present modeling relies on the parcellation of brain regions. As such, a delicately charted brain map^55^ would be preferred to provide executable guidance with greater precision, although it heightens the risk of increased model complexity and computation load. Thus, more insights into the biological validity of the present model using well-controlled animal studies ^49^ will help to balance this tradeoff.

Taken together, these findings, although exploratory in nature and requiring clinical validation, advance an unprecedented view of the global dynamics of brain function through selective manipulation of a local brain area. This suggests that the ability to simulate, predict and assess perturbation of large-scale network dynamics evoked by focal stimulation may unfold a new dimension in the increasingly attractive field of brain connectomics, with which one can describe the dynamic spread and functional consequences of pathological processes. The development of neural circuit-based guidance approaches, coupling whole-brain connectomic modeling to clinical considerations of therapeutic intervention plans, which is yet to be established, may help to reach this goal and possibly diminish the risk of the DBS-linked side effects. Stratification by patient-specific functional connectome may have predictive value in respect to the efficacy of a neuromodulation treatment and even guide its choice, providing a stepping-stone in the advancement of translational progress in precision medicine.

## Methods

### Participants

Demographic and clinical characteristics of 179 participants used in this study are summarized in Table 1. The locally enrolled sample consisted of 94 subjects, among whom 47 were patients recruited at the Department of Neurology at Ruijin Hospital with a diagnosis of PD according to UK Brain Bank criteria. All these PD patients received medication and the severity of motor symptom in patients was evaluated by study-site personnel and qualified neurologists using the Unified Parkinson’s Disease Rating Scale (UPDRS-III) and the Hoehn-Yahr Staging Scale. 47 healthy comparison (HC) subjects were recruited by advertisement and administered the Mini-International Neuropsychiatric Interview (MINI 6.0.0). All recruited subjects were without any neurological or psychiatric condition, a history of substance abuse, brain injury or other notable abnormality upon MRI examination. Four PD patients and one healthy subject were excluded due to severe head motion during MRI scanning (larger than 2 mm translation displacement or 2.0° rotation). This study was approved by the Institutional Review Board at Ruijin Hospital of Shanghai Jiao Tong University and by the Biomedical Research Ethics Committee, Shanghai Institutes for Biological Sciences, Chinese Academy of Sciences. Informed consent was obtained from each subject or their legal guardian upon receiving a complete description of the study.

**Table 1.** Demographic and clinical characteristics of local PD patients, public PD patients and healthy comparison subjects.

|  | Local PD<br>(N = 43) | Public PD<br>(N = 90) | HC<br>(N = 46) |
| --- | --- | --- | --- |
| Age (years) | 62.00 ± 6.04 | 61.70 ± 10.33 | 31.37 ± 8.30 |
| Gender (No. of males/females) | 21/22 | 61/29 | 27/19 |
| Duration of illness (years) | 4.41 ± 2.54 | 2.24 ± 1.04 | - |
| UPDRS-III | 21.93± 11.39 | - | - |
| MDS-UPDRS-III | - | 21.28 ± 11.00 | - |
| H-Y scale | 1.62 ± 0.53 | 1.73 ± 0.47 | - |

Values are mean ± SD. UPDRS-III, Unified Parkinson’s Disease Rating Scale part III; MDS, Movement Disorder Society; H-Y scale, Hoehn and Yahr scale.

Data for an additional set of 93 PD patients with resting-state fMRI were obtained from the Parkinson’s Progression Markers Initiative (PPMI) database. Three cases were excluded due to severe head motion using the same exclusion criteria. All data are fully anonymized as required by HIPAA regulations and participating sites received local Institutional Review Board approval for acquisition of contributed data. Detailed information about PPMI patients is available at (www.ppmi-info.org).

### MRI data sets

All local participants were scanned with a standard 12-channel head coil on a Siemens Tim Trio 3.0 T scanner (Erlangen, Germany), located at the Institute of Neuroscience, Chinese Academy of Sciences. Each participant was instructed to lie supine in the scanner wearing earplugs, with their head snugly fixed by tight but comfortable foam pads. Real-time electrocardiogram output and heart rate were continuously monitored throughout the scan (Erlangen, Germany). High-resolution T1-weighted images were acquired using a 3D magnetization-prepared rapid gradient-echo sequence (TR/TE = 2300/3 ms, TI = 1000 ms, flip angle = 9°, FOV= 256×256 mm^2^, voxel size = 1×1×1 mm^3^, 176 consecutive sagittal slices). Resting-state fMRI images for patients were acquired for 300 volumes using a multiband echo planar imaging sequence (TR/TE = 2000/30ms, flip angle = 62°, in-plane resolution = 3×3 mm^2^, slice thickness = 3 mm, 56 axial slices, multiband acceleration factor = 2). Note that local PD patients had received their medication accordingly at the time of the scan. For healthy subjects, resting-state fMRI images were acquired for 300 volumes with a single-shot echo planar imaging sequence (TR/TE = 3000/30ms, flip angle = 90°, in-plane resolution = 2×2 mm^2^, 3 mm slice thickness, 47 axial slices). During the resting-state scan, participants were instructed to lie still in the scanner with their eyes closed, remain awake, and not think of anything in particular (adherence was confirmed by participants immediately after the scan).

Anatomical and resting-state fMRI data from the public PPMI dataset were acquired on Siemens Tim Trio 3.0 T scanners. Structural images were recorded using a similar protocol as described above. Resting-state fMRI images were acquired for 210 volumes using an echo-planar imaging sequence (TR/TE = 2400/25 ms, flip angle = 80°, in-plane resolution = 3.3×3.3 mm^2^, slice thickness = 3.3 mm, 40 axial slices). Further technical details can be found in the MRI operation manual available at http://www.ppmi-info.org/.

### Network construction

The fMRI data were minimally preprocessed using Statistical Parametric Mapping (SPM, http://www.fil.ion.ucl.ac.uk/spm). The first 10 volumes were discarded for signal equilibrium and the remaining volumes were corrected for temporal difference in slice acquisition and rigid-body head movement. The corrected data were spatially normalized to the MNI (Montreal Neurological Institute) space and resampled to 3 mm isotropic voxels. After normalization, six motion parameters (three for translation and three for rotation) estimated during the realignment process were regressed out and linear drift was removed. A band-pass filter (0.01 – 0.08 Hz) was applied to remove the low frequency drift and high frequency respiratory and cardiac noise. We constructed the whole-brain connectivity network using a custom parcellation scheme based on the standard Automated Anatomical Labeling (AAL) atlas with the addition of the subthalamic nucleus as defined by the ATAG subcortical atlas ^56^ (the details of a total of 92 brain regions are listed in Supplemental information **Table S1**). The time series of all voxels within each region were extracted and averaged to obtain a mean time series. Functional connectivity *f*_*ij*_ between brain regions was represented by calculating the Pearson correlation coefficients between the mean time series of any pair of parcellated regions (*i* and *j*). A 92×92 connectivity matrix **F** was generated for each subject and then subjected to Fisher’s Z-transformation for subsequent analysis.

### Whole-brain network model of neurostimulation

We first simulated the network effects of neurostimulation by considering the constructed connectome **F** as the resultant network of information spread or diffusion over a direct network **D**. The rationale here is that the local effect is the action of stimulation imposed on the direct network **D**, and the globally distributed effect is the propagation result of the local effect on the direct network. As such, we adopted a network deconvolution algorithm ^57^ to derive a direct network **D** from the observed network **F**, and used the corresponding transformation rule, transitive closure, to simulate the global effects of local stimulation on a large-scale neuronal network. A detailed description of the modeling is provided in the Supplemental Information. We then denote a hypothetical neurostimulation event as a regulation operator **P**, which is imposed onto the direct network **D** (i.e.,**D**′ = **D**⊙**P**, where ⊙ is the element-wise product of two matrices). If neurostimulation targets one pair of bilateral loci, most **P** entries will have a value of one, with the exception of two columns and two rows that are directly connected to the locus. For simplicity, the same value that represents the neurostimulation strength is applied to the remaining entries in **P**: a value larger than one signifies up-regulation, whereas a value smaller than one indicates down-regulation (e.g., 1.5 represents a 50% up-regulation, 0.7 represents a 30% down-regulation). Finally, the neurostimulation-tuned direct network **D**′ is convolved with transitive closure to obtain the post-neurostimulation connectivity matrix **F**′ (Figure 1A).

The simulated outcome of neurostimulation was evaluated by quantitatively measuring the similarity between a group-averaged healthy matrix obtained from healthy subjects and an individual or group-averaged post-neurostimulation matrix obtained from PD patients. Connectomic similarity was quantified as the Pearson correlation coefficient between two vectorized (concatenated by rows) upper triangles of the connectivity matrix. This index was appropriate to characterize the discriminant difference between HC and PD patients, as demonstrated in Figure S6A and S6B. In order to compare the outcomes across brain regions, individuals, and groups, we standardized all outcome similarities using *relative change*, defined as follows:

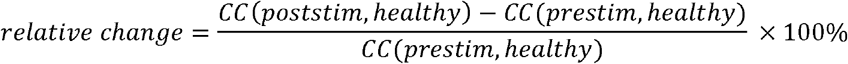

Relative change was used as a quantitative measure to represent the percentage improvement in connectomic similarity towards a healthy regime that is achieved by neurostimulation. The higher the connectomic similarity with the matrix of the healthy subject group, the better the simulated therapeutic effect is. At each targeted brain region, the up- or down-regulation magnitude that resulted in the highest *relative change* was deemed as the best strength. All target regions were then ranked according to their *relative change* at optimal neurostimulation strength. Higher ranks indicate desirable stimulation candidates with better therapeutic effects. All predictions were made at both single patient and patient group levels, as illustrated in Figure 1B.

### Study design, statistical analysis and cross-validation

As a proof-of-principle study, we rigorously tested the robustness and consistency of the predicted results using various cross-validation procedures. To validate whether the model is biased by the inclusion or exclusion of subject data, we randomly sampled half of the PD and HC groups 1000 times for predictions. Unblinding of between-group labels is also necessary to generate cross-validated prediction accuracies. We randomly sampled half of the HC group 1000 times as the “patient group” to validate whether the present strategy for target prediction is biased by the modeling *per se*. Moreover, we generated a new 1024-region parcellation template to examine whether the prediction of targets or strengths was biased by a specific brain parcellation scheme. As for the results of individual patients, we assessed the statistical significance (one sample t-test) and the effect size (Cohen’s *d* value) of the simulated therapeutic effects for all brain regions. We assigned the rank for each brain region across individuals and summarized the occurring times of the ranks. The optimal neurostimulation strength of each region for each individual patient was also plotted and summarized statistically (one sample t-test). For external validation, we repeated the same procedure using an independent public dataset (PPMI PD patients).

To reveal the extent of rectification of network topology achieved through neurostimulation, we conducted edge-wise statistical comparison between the two groups. The results of two sample t-tests after Bonferroni correction for multiple comparisons (p < 0.05 corrected) were categorized into three types: removed abnormal connections (i.e., significantly different between the PD and HC groups before, but not after neurostimulation); newly emerged abnormal connections, and unchanged abnormal connections. Finally, to examine the relationship between the predicted priority of brain regions and symptom severity, a Kendall rank correlation was calculated between each target’s rank and UPDRS-III score for two datasets.

## Acknowledgments

This work was supported by the Hundred Talent Program of the Chinese Academy of Sciences (Technology), the Strategic Priority Research Program (B) of the Chinese Academy of Sciences (XDB02050006) (Z.W.), the National Natural Science Foundation of China (81571300, 81527901, 31771174 to Z.W.; 81271518, 81471387 to B.M.S.), and National Key R&D Program of China (2017YFC1310400 to Z.W.). Part of data used in the preparation of this article were downloaded from the Parkinson’s Progression Markers Initiative (PPMI) database (www.ppmi-info.org/data). For up-to-date information on the study, visit www.ppmi-info.org. PPMI, a public-private partnership, is funded by the Michael J. Fox Foundation for Parkinson’s Research and funding partners, including Abbott, Avid Radiopharmaceuticals, Biogen Idec, Bristol-Myers Squibb, Covance, Ian, GE Healthcare, Genentech, GSK-GlaxoSmithKline, Lundbeck, Lilly, Merck, MSD-Meso Scale Discovery, Pfizer, Piramal, Roche, Servier, and UCB (www.ppmi-info.org/fundingpartners). The authors thank Drs. Mu-ming Poo and Wayne Goodman for their insightful comments on the manuscript and related topics.

## Author contributions

Z.W. and B.M.S. were responsible for the concept and the design of the study. X.Y.C., C.C.Z., Y.X.L., W.W.Y., P.H., and S.D.C. acquired the data. X.Y.C., C.C.Z., Y.X.L., Q.L., and P.H. analyzed all or parts of the data. X.Y.C., C.C.Z., Y.X.L., and Z.W.
drafted all or parts of the article, with input from other authors. Z.W., B.M.S., and
S.D.C. obtained the funding and supervised the study.

**Supplementary Information** accompanies this paper available at online.

